# RoAM: computational reconstruction of ancient methylomes and identification of differentially methylated regions

**DOI:** 10.1101/2024.08.08.607143

**Authors:** Yoav Mathov, Naomi Rosen, Chen Leibson, Eran Meshorer, Benjamin Yakir, Liran Carmel

## Abstract

Identifying evolutionary changes in DNA methylation bears a huge potential for unraveling adaptations that have occurred in modern humans. Over the past decade, computational methods to reconstruct DNA methylation patterns from ancient DNA sequences have been developed, allowing for the exploration of DNA methylation changes during the past hundreds of thousands of years of human evolution. Here, we introduce a new version of RoAM (Reconstruction of Ancient Methylation), a flexible tool that allows for the reconstruction of ancient methylomes, as well as the identification of differentially methylated regions between ancient populations. RoAM incorporates a series of filtering and quality control steps, resulting in highly reliable DNA methylation maps that exhibit similar characteristics to modern maps. To showcase RoAM’s capabilities, we used it to compare ancient methylation patterns between pre- and post-Neolithic revolution samples from the Balkans. Differentially methylated regions separating these populations are shown to be associated with genes related to regulation of sugar metabolism. Notably, we provide evidence for overexpression of the gene PTPRN2 in post-Neolithic revolution samples. PTPRN2 is a key regulator of insulin secretion, and our finding is compatible with hypoinsulinism in pre-Neolithic revolution hunter-gatherers. Additionally, we observe methylation changes in the genes EIF2AK4 and SLC2A5, which provide further evidence to metabolic adaptations to a changing diet during the Neolithic transition. RoAM offers powerful algorithms that position it as a key asset for researchers seeking to identify evolutionary regulatory changes through the lens of paleoepigenetics.

## Introduction

Identifying changes in gene expression levels is a powerful tool to study evolutionary shifts and adaptations. Specifically, many efforts have been directed to identifying changes in gene regulation along the human lineage (1–3). The rise of ancient DNA (aDNA) offers a potential to study regulatory changes that shaped modern humans and might have affected our recent evolution. Given that changes in gene regulation are difficult to read directly from the DNA sequence, and that RNA rarely survives in ancient remains, DNA methylation stands out as the best proxy of ancient gene expression levels (2, 4). DNA methylation, which in mammals affects cytosines in the context of CpG positions, is a key epigenetic mark that is tightly associated with gene expression levels (5). Incidentally, unlike other epigenetic marks such as histone modifications, DNA methylation is highly stable and remains on aDNA for extended periods (6).

More than a decade ago, we and others developed computational methods for reconstructing premortem DNA methylation patterns from aDNA (7, 8). These techniques used the fact that deamination – the main chemical degradation of aDNA – turns methylated cytosines into thymines and unmethylated cytosines into uracils (6). In preparing the aDNA for sequencing, a treatment with uracil-specific excision reagent (USER) is often used (6), generating an asymmetry between methylated and unmethylated cytosines, which can be used to distinguish between them by carrying out statistical analysis of the number of *C* → *T* transitions in CpG positions. A high rate of *C* → *T* indicates premortem hypermethylation, while the converse indicates hypomethylation. These pioneering methods have not only opened avenues for exploring epigenetic regulations in ancient samples, but also enabled the reconstruction of genome-wide methylation maps and the identification of differentially methylated regions (DMRs) between ancient samples. These works founded the field of paleoepigenetics, which provided significant insights into various aspects of human evolution (2, 4, 9)

Several of these aDNA reconstruction algorithms were published as tools, including RoAM (8), epiPALEOMIX (10), and its successor DAMMET (11). However, RoAM was distributed as Matlab code, and its use was limited. DAMMET, which employs a maximum likelihood estimation to calculate methylation levels based on the *C* → *T* transition counts, is the most recent tool, but has several limitations. First, it is assumed that all four nucleotides have the same frequency of 0.25, ignoring GC content biases. Specifically, the GC content of the human genome is known to be 40.9% (12). Second, it is assumed that there is an equal probability of 1/7 for each dinucleotide that can be read as a CpG due to a mutation, which does not align with established mutation rates in humans (13, 14). Notably, mutations are not uniformly distributed, with *C* → *T* mutations being particularly prevalent (15). Third, the estimation procedure includes cytosines outside of a CpG context, which are assumed to be unmethylated. However, some small levels of non-CpG methylation are known to exist (16), especially in embryonic cells and specific mature cell types such as neurons. Not much is known about the prevalence and significance of non-CpG methylation in tissues that are present in the fossil record, particularly bones and teeth. This, combined with the assumption that the deamination rate of cytosines in non-CpG context is identical to that of cytosines within CpG context, can introduce bias into the reconstructed map. Indeed, previous works showed that methylation maps provided by DAMMET show more hypomethylation than expected (17, 18).

Moreover, DAMMET generates methylation maps, but does not compare them to identify DMRs. RoAM, on the other hand, does include a method to detect DMRs, but could originally do so only between pairs of samples. As the number of published aDNA samples continues to grow, the detection of DMRs between large groups of samples has become desirable, as they have the potential to unveil DNA methylation differences between populations, within the same population across different time points, and between closely related species.

Here, we present a new python version of RoAM (Reconstruction of Ancient Methylation), which removes many of the aforementioned limitations. It is feature-rich, flexible, easy to use, and its code is freely available. It allows for the generation of premortem genome-wide aDNA methylation maps, as well as the detection of DMRs between groups of samples. The current version contains numerous improvements over the original software, including novel methods for filtering out true *C* → *T* mutations. RoAM does not use assumptions about nucleotide frequencies, and does not rely on cytosines outside of CpG context.

To demonstrate the capabilities of RoAM, we reconstructed the methylomes of 14 samples from the Balkans and detected DMRs between pre-Neolithic revolution specimens and post-Neolithic revolution ones. As expected from the short time span separating these populations, we detected only four DMRs with modest levels of methylation change. However, the genes that are associated with these DMRs might hold clues to nutritional changes that occurred during the Neolithic transition. Notably, PTPRN2, which is involved in insulin response to glucose stimulus, is predicted to be overexpressed in post-Neolithic revolution individuals, as expected for high carbohydrate diet. Methylation changes were also found in EIF2AK4, a sensor for amino acid deprivation that also regulates insulin, and SLC2A5, the main fructose transporter. In total, these findings may provide clues to regulatory changes that might have accompanied the major changes in diet and lifestyle that ensued following the Neolithic revolution.

## Methods

RoAM performs two main tasks. First, it reconstructs the methylation map of ancient samples. Then, it compares groups of samples to detect DMRs between them (Figure 1). In the following, we thoroughly describe these two parts.

**Figure 1.**
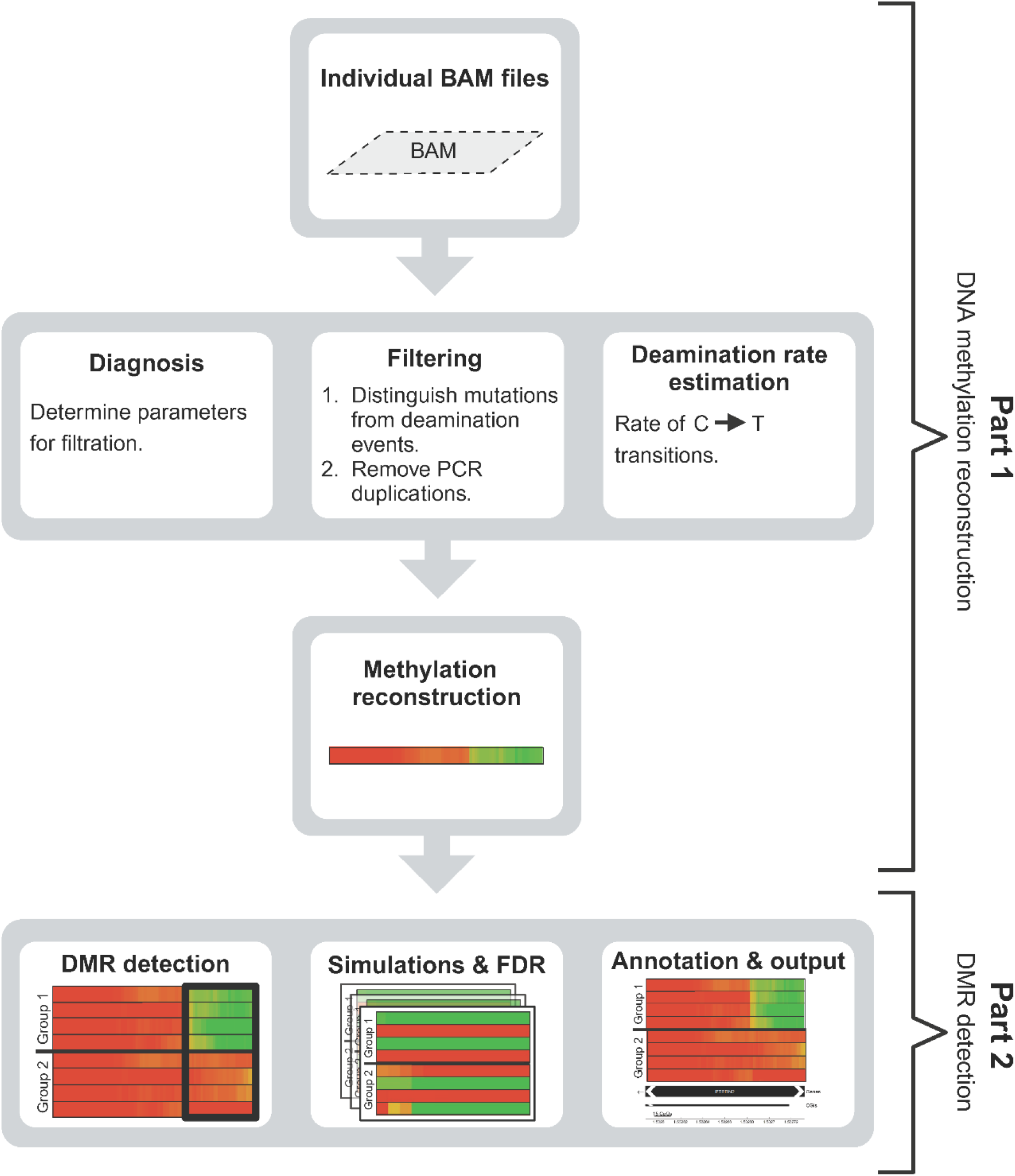
The RoAM pipeline is split into two parts. In Part I, RoAM starts with BAM files of ancient genomes, and reconstructs the individual methylation maps. In Part II, RoAM detects differentially methylated regions (DMRs) distinguishing two groups of ancient samples. (created with Biorender)

### Part I. Reconstructing premortem ancient DNA methylation

The main input for this part is a BAM file of an ancient individual. Each BAM file is analyzed through five consecutive steps (Figure 1): (1) basic processing of the BAM file; (2) diagnosis step to automatically determine filtering parameters; (3) carrying out the filtering to remove non-informative CpG positions; (4) estimating the deamination rate; and (5) reconstructing the premortem DNA methylation.

In addition to the BAM file, RoAM requires several input parameters that instruct it how to process the specific sample. Although not mandatory, it is highly recommended to provide a reference present-day DNA methylation map generated from the same tissue from which the aDNA was extracted. The reason for this is that such a reference allows for more accurate estimation of the deamination rate and methylation reconstruction. We provide such reference for present-day human bone aligned to hg19, see https://carmelab.huji.ac.il/data.html. A full description of all input parameters can be found in the README file in the GitHub page of RoAM, https://github.com/swidler/roam-python.

RoAM provides the user with two outputs that contain the reconstructed methylation map. One is a simple BED file, and the other is a python object that also contains all parameters that were used by the algorithm. This python object is later required for the DMR detection part.

#### Step I: BAM file processing

RoAM reads all relevant information from a BAM file into a python object. This object stores some descriptive characteristics of the sample, such as the sample name, species and library preparation method. In addition, it holds relevant summary statistics of the sequencing data, specifically nucleotide counts in each CpG position. At later steps, more information is stored in this object, such as the filtering parameters, the estimated deamination rate, and the reconstructed methylation values. BAM files are processed, one chromosome at a time, using Python’s *pysam* module (https://github.com/pysam-developers/pysam). During this step, RoAM filters out low quality reads, and computes base counts for each position in a way that depends on the library preparation method. For single-strand libraries, each strand is treated independently, and only *C* → *T* transitions are relevant. For double-strand libraries, complementary *G* → *A* events along the opposite strand are counted as well.

This step is the most time-consuming part of the pipeline. However, it needs to be executed only once per sample, after which the saved object can be re-used.

#### Step II: Diagnosis

Filtering in RoAM is aimed at removing CpG positions whose cytosine and thymine counts may be spurious or uninformative to the reconstruction process. We identify two groups of such CpG positions. First, some positions show suspiciously high coverage that suggest they are a result of PCR duplications. Second, some positions show *C* → *T* counts suggestive of premortem mutations rather than deamination.

RoAM runs a diagnostic procedure that automatically suggests parameter values that should be used during filtering. The user can manually override these suggestions.

##### Identifying PCR duplications

Let *t*_*i*_ and *c*_*i*_ be the counts of thymines and cytosines, respectively, in CpG position *i*, and let *n*_*i*_ = *t*_*i*_ + *c*_*i*_ be the total count in this position. As a first step, we use a crude outlier removal process to remove positions with extreme values of *n*_*i*_. Then, we use a more refined method to remove additional outlying positions. For the first step, we compute the 25^th^ and 75^th^ percentiles of all *n*_*i*_ values, denoted *p*_25_ and *p*_75_, respectively. We compute the interquartile range as *r* = *p*_75_ − *p*_25_, and remove all positions where *n*_*i*_ > *p*_75_ + *s* · *r*. Here, *s* is a parameter called *span*, set by default to 5.

For the second step, let *N*(*c*) be the histogram of the remaining *n*_*i*_ values, counting the number of CpG positions with coverage *c. N*(*c*) resembles normal distribution truncated at 1, and with a heavier right-tail. To account for the truncation, we use the following method to estimate the parameters of the distribution. Let *c*_*m*_ be the coverage level that maximizes *N*(*c*). Based on the three coverage levels *c*_*m*_ − 1, *c*_*m*_, and *c*_*m*_ + 1, we estimate the parameters *a, b*, and *c* of the best-fitting binomial *N*(*x*) = *ax*^2^ + *bx* + *c*. We estimate *μ*, the mean of the normal distribution, as the point where this function is maximized, 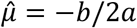. We call the maximum value of the function 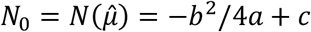. To estimate the standard deviation, *σ*, we find the first bin in the histogram, *b*, such that *b* is greater than *μ* and for which *N*(*b*) ≤ *t* · *N*_0_, where *t* is a parameter that is set by default to 0.1. The smaller *t* is, the more the approximation accounts for the heavy tail. Let *f* = *N*(*b*)*/N*_0_ be the ratio of these two bins. Then,

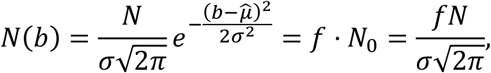

where *N* is the total number of counts, *N* = ∑_*c*_ *N*(*c*). Therefore, *σ* can be estimated using this ratio by

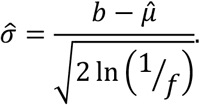

Given the estimates 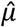 and 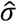, we further remove CpG positions that might be a result of PCR duplicates by filtering out all positions whose coverage exceeds a threshold *c*_*T*_, determined as the coverage that would yield an expected value of counts less than one. We find *c*_*T*_ by solving for

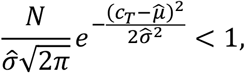

giving

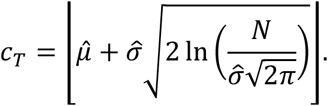

Removal of PCR duplicates is done by removing all positions with

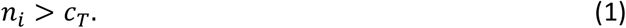

##### Detecting C → T mutations

*C* → *T* mutations are detected in two ways. As we detailed in past papers (8, 19), in aDNA that was sequenced using single-strand libraries, mutations can be distinguished from deamination by examining the opposite strand for *G* → *A* transitions.

Here, we developed an alternative technique, that is also applicable for aDNA sequenced using double-strand libraries. It is based on the analysis of the histogram *H* of *t*_*i*_ and removing CpG positions with *C* → *T* rates that are too high and likely represent a *C* → *T* mutation. We examine each coverage level *c*, independently, and therefore can now assume that we are only looking at the *N*(*c*) CpG positions whose coverage is *c*. Denoting by *p* the probability of a *C* → *T* transition, we assume that *H* represents a mixture of three binomial distributions: (1) homozygous mutations, where *p*_1_ ≈ 1. (2) heterozygous mutations, where *p*_2_ = 0.5. (3) non-mutated sites that went through deamination, where *p*_3_ ≈ 0.01; We use expectation maximization (EM) to estimate the *p*’s and *w* ‘s in the binomial mixture model (BMM):

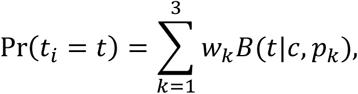

where *w*_*k*_ is the weight of the *k*’th distribution. According to the EM formulation, in each iteration we update the following three magnitudes:

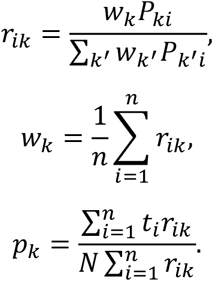

Here, *P*_*ki*_ = *B*(*t*_*i*_|*c, p*_*k*_). To make the computations more efficient, we avoided summing over *i* = 1, …, *n* and rather summed over the values of the histogram, *g* = 0, …, *N*. Hence, we modified the above iterations to read:

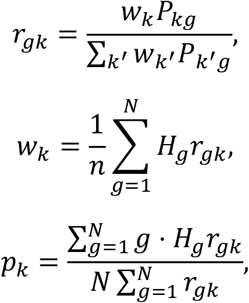

where *P*_*kg*_ is the value of the *k*’th distribution for the value of the *g*th bin. Once we have estimated all these parameters, we decide on the threshold *k*_*c*_, defined as

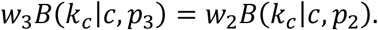

Writing the binomial distribution explicitly, we get

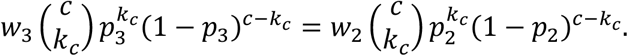

Taking log from both sides,

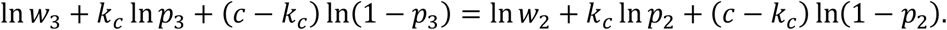

This gives

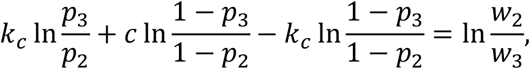

or

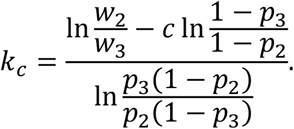

In our case, we force *p*_2_ = 0.5, hence

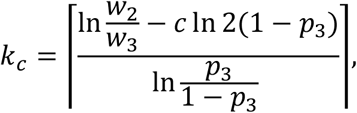

where the ⌈·⌉ operator means the closest integer from above. We then remove all CpG positions whose coverage is *c* and where *t*_*i*_ ≥ *k*_*c*_.

We can evaluate the number of true deaminated positions that are missed by this filtering,

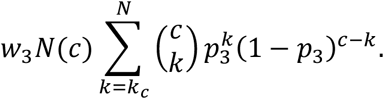

Similarly, we can evaluate the number of positions with false deamination, which is

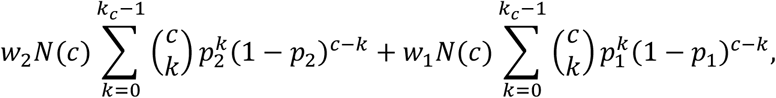

which, after substituting *p*_2_ = 0.5, gives

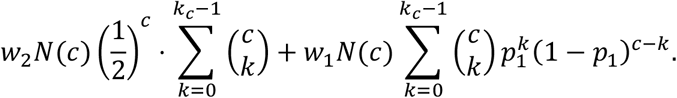

These computations of false negative and false positive rates are reported during the diagnosis step. Next, from within all positions with coverage 1 ≤ *c* ≤ *c*_*T*_, we remove those for which

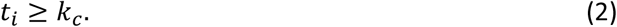

#### Step III: Filtering

The filters outlined in Step II are used to determine which sites to remove. First, we clean PCR duplicates. This is done by setting a threshold *c*_*T*_ and removing every site where *n*_*i*_ > *c*_*T*_ (see Eq. 1). Then, let *i* denote all the sites with a given coverage level *c*, where *c* = 1, …, *c*_*T*_. We remove all sites for which *t*_*i*_ ≥ *k*_*C*_ (see Eq. 2), as they are suspected as true mutations.

For single-strand libraries, we add a filter on the *G* → *A* transitions in the opposite strand. To this end, we set a maximum number of allowed A’s, *a*_*L*_, as well as a minimum *G* → *A* ratio, *T*_*m*_ (default: 0.25). Then, we remove all sites where

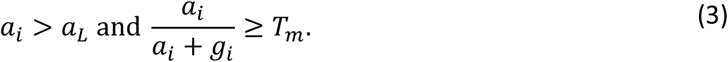

After filtering, a merging procedure is applied to combine information from both Cs of the same CpG position (on opposite strands). The methylation state of these two Cs should be identical (20), thus merging the counts of Cs and Ts from both strands increases the amount of information obtained from each CpG position.

#### Step IV: Estimation of deamination rate

The deamination rate is estimated using the same technique we have previously detailed (19), and is based on the C and T counts in CpG positions whose methylation in the modern reference is above a certain threshold, *m*_*h*_. This parameter should be close to one and is exactly one by default. As the maximum-likelihood estimator of the methylation in a site is

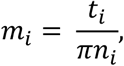

limiting ourselves to positions where *m*_*i*_ = 1 lets us estimate the degradation rate by

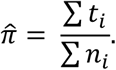

In a case where a reference is not available, one can estimate the degradation rate by assuming knowledge of the global mean methylation in the sample, *m*_*g*_. Then, we estimate the deamination rate by

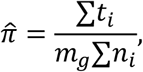

where the sum is over all positions in the genomes.

This function also computes the local methylation rate for each chromosome, which allows testing for homogeneity of the deamination rate across chromosomes.

#### Step V: Methylation reconstruction

By default, RoAM uses histogram matching to reconstruct the methylation maps by finding the non-linear transformation *m*_*i*_ = *f*(*t*_*i*_ */n*_*i*_) that makes the histogram of *m*_*i*_ as close as possible to that of a reference methylation map in a modern bone (21, 22), where *m*_*i*_ is the estimated methylation at position *i*.

By design, we obtain a histogram of methylation values which resembles the equivalent histogram from modern-day sample. It stands in contrast to DAMMET, which tends to show shifts towards low methylation (Figure 2). We used the *χ*^2^ statistic to compare the histograms of maps reconstructed using RoAM and DAMMET to a bone map that was not used as a reference. Whereas 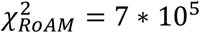, the same statistic for DAMMET was one order of magnitude larger, 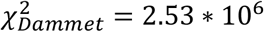, indicating larger distance between the histograms.

**Figure 2.**
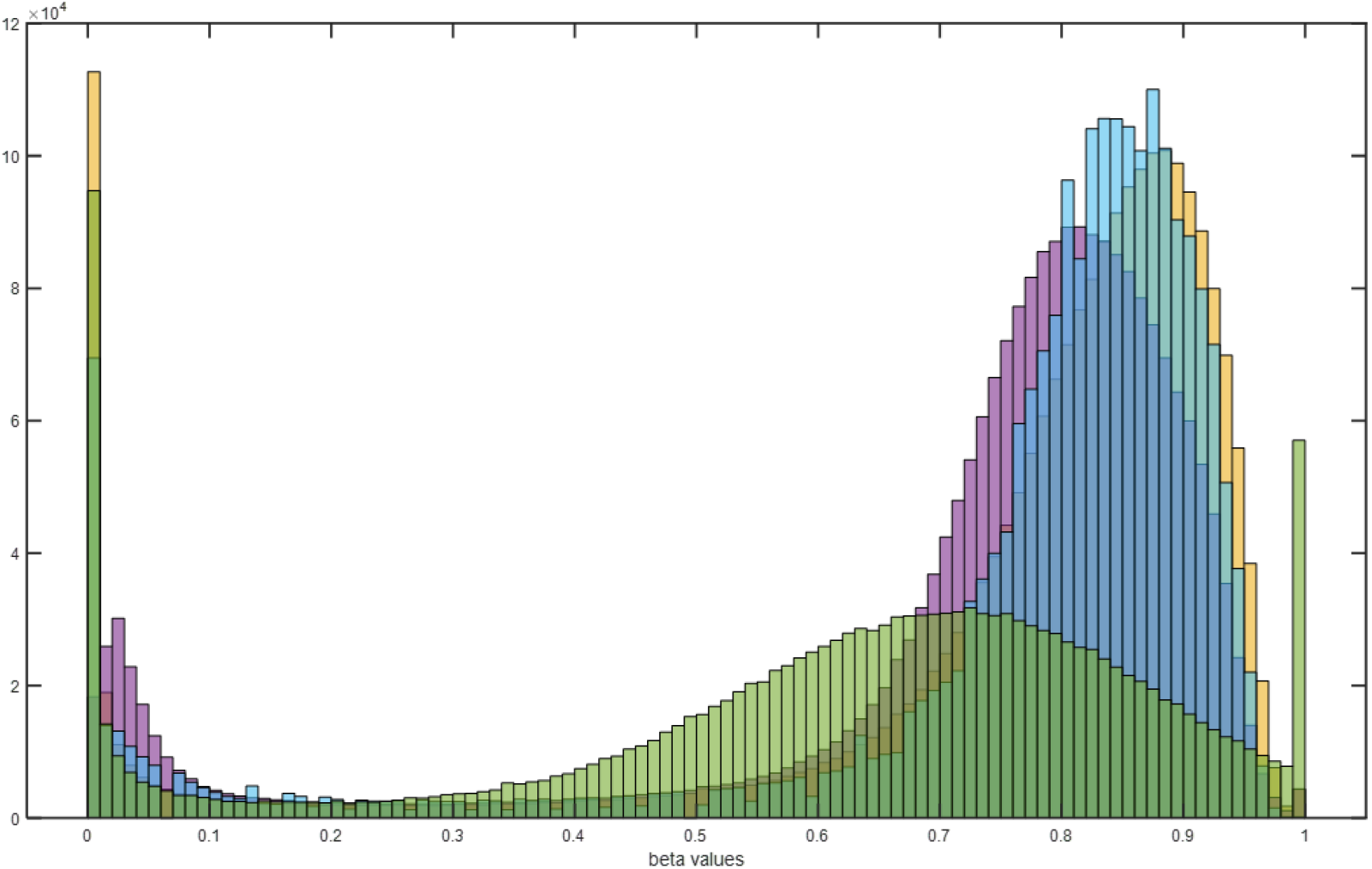
Genome-wide histograms of methylation levels (chromosome 1). Ancient DNA methylation in sample I1116 was reconstructed by RoAM (blue) and by DAMMET (green). Two modern bone samples are shown. One (yellow) was used as a reference in RoAM, and another (purple) that was not used in RoAM.

Histogram matching serves as the default method for methylation reconstruction, but users may choose two other methods. One is the truncated linear transformation,

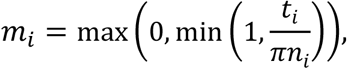

as described and used previously (19). To achieve a smooth truncation, RoAM also offers a third method, called the logistic transformation, where

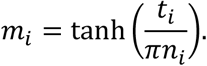

Given the typically small thymine counts in each CpG position, *t*_*i*_, RoAM reconstructs methylation in windows of *W* consecutive CpG positions (*W* is always set as an odd number, and the reconstructed methylation in the window is assigned to the middle position). The user may determine the window size they wish to use, but RoAM includes two methods to automatically determine the window size.

##### Probability-based method

We require that the probability of observing no thymines in a window for a minimum methylation level *m*_0_ be less than *p*_0_. This translates into

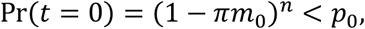

where *t* is the total thymine count in the window, and *n* is the total count of thymines and cytosines. Taking log of both sides we get

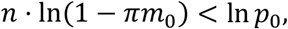

meaning that we have to have

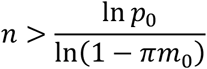

in the window. If the window is covered by the average effective coverage, *C*, then *n* = *WC*. This translates into the following window size:

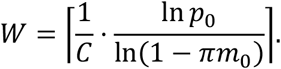

##### Relative-error-based method

We require that the relative error in estimating the methylation, when the true methylation is *m*_0_, be lower than 1*/k*. Let the estimator for the methylation be

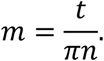

Then, its mean is

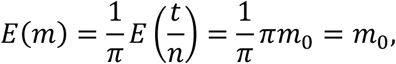

and its variance is

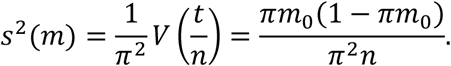

We require that *s*(*m*)*/E*(*m*) < 1*/k*, hence

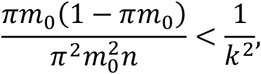

or

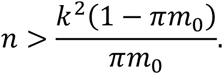

Again, using the average effective coverage, *C*, this translates into

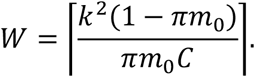

The default that RoAM uses is the probability-based method.

### Part II. DMR detection

Once reconstruction of methylation has been achieved for multiple samples, a second part of RoAM is designed to detect and statistically validate DMRs between two groups of samples. This process comprises the following steps (Figure 1): (1) DMR detection between the two groups, (2) the use of simulations to adjust the parameters of the DMR-calling algorithm to reach a desired level of false discovery rate (FDR), and (3) annotation of the final list of DMRs. The algorithm provides a table with a list of all the DMRs, their location, annotation, and the methylation level in each of the samples, as well as the combined estimated methylation in each group.

#### Step I: DMR detection

Let us first examine a group of *S* samples. We assume that the methylation across members of the group is homogeneous, and denote the common methylation value in window *j* as *m*_*j*_.

Let us look at sample *i*. We assume that the observed number of T bases in window *j* is binomially distributed, *t*_*ij*_ ∼*B*(*n*_*ij*_, *m*_*j*_*π*_*i*_), where *π*_*i*_ is the deamination rate of the sample and *n*_*ij*_ is the sum of the Cs and Ts in each CpG in window *j* in sample *i*. The likelihood of sample *i* is

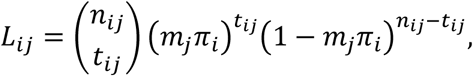

and the log-likelihood

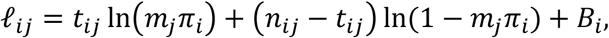

where *B*_*i*_ is a term that is independent of *m*_*j*_. The total log-likelihood of all *S* samples in the group

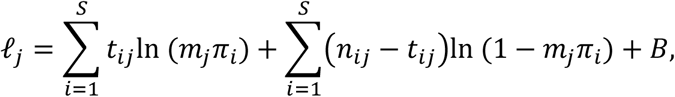

where *B* = ∑_*i*_ *B*_*i*_ is a term independent of *m*_*j*_. The score function with respect to *m*_*j*_ is:

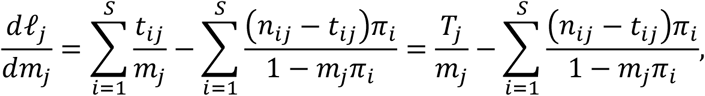

where

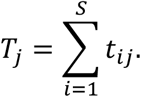

To look for the maximum likelihood estimator, we should equate this to zero. This can be done numerically, using, e.g., the Newton-Raphson method. For this,

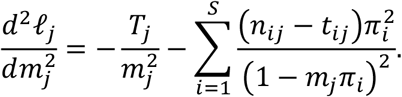

Given 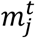 is the approximate solution at iteration *t*, the solution at iteration *t* + 1 is given by

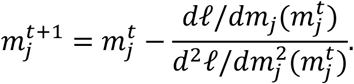

In order to get an initial guess, we may obtain an approximated solution using

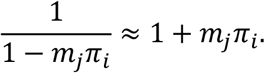

Hence,

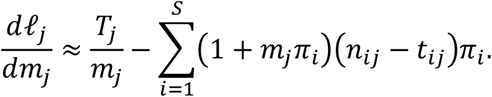

This simplifies into

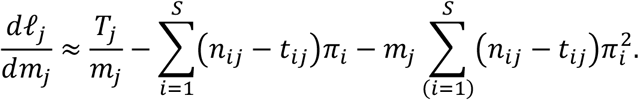

Further approximating by neglecting terms of the order of 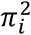, we get

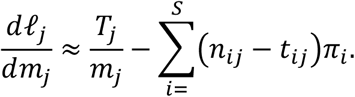

This can be written as

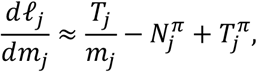

where

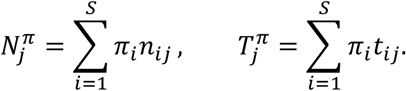

The approximate solution is therefore

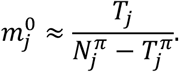

The Fisher information for estimating *m*_*j*_ is equal to the expectation of the negative second derivative of the log-likelihood function,

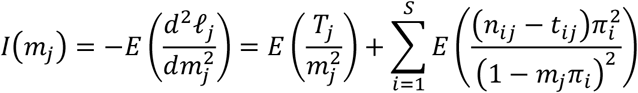

The empirical Fisher information is the evaluation of this negative second derivative at the estimated value of the parameter, and may serve as an approximation of the Fisher information,

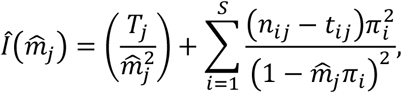

where 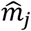 is the estimator obtained from the iterations of the Newton-Raphson algorithm. The empirical Fisher information is computed as part of the implementation of the algorithm. Finally, we may approximate the variance of the estimator via the inverse of the empirical Fisher information:

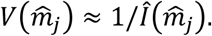

After computing *m*_1,*j*_ and *m*_2,*j*_, the methylation levels in every genomic window for the two groups, we can compare the two using a similar approach to the one used in (19). To this end, we define the two statistics

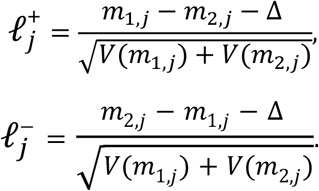

Here, Δ is a parameter of the algorithm, associated with the desired minimal methylation difference we wish to detect between the two groups. Next, we use a cumulative sum for 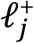 and 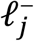 to identify DMRs, as described in (19). In brief, we define the vectors *Q*^+^ and *Q*^−^ of the same length as 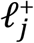 and 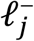, as

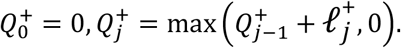

*Q*^−^ is defined in an analogous way.

In DMRs where group 1 is hypermethylated compared to group 2, we will obtain a sequence of positive values for *ℓ*^+^ resulting in an elevation in the values of *Q*^+^. We define the DMR as the region [*a, b*] between the last zero *Q*_*a*_ = 0, and the highest value *Q*_*b*_ = *Q*_*max*_ up to the next zero (19).

Each DMR is characterized by several properties, such as its genomic length, the number of CpG positions it harbors, and its *Q*_*max*_. This allows for further filtering of DMRs, to achieve a desired false discovery rate (FDR), as explained in the next section.

#### Step II: Simulations and FDR

We employ simulations to filter out DMRs in such a way that we achieve a desired FDR level. The detailed procedure can be found in our previous paper (19). In short, we imitate the deamination process in each sample by generating Ts using a binomial process, where the coverage in each position and the deamination rate are kept constant for each sample. The methylation value in each position is determined in advance, and is kept constant across the samples from both groups, to model the null hypothesis of no methylation differences between the groups.

Subsequently, we apply the same DMR detection procedure to the simulated data and count the number of detected DMRs. This number represents the number of DMRs detected under the null hypothesis. Repeating this many times (typically 100 times), we may compute the expected fraction of false DMRs within our original list of DMRs. By default, we set an FDR threshold of 0.05, but this parameter can be adjusted by the user. Given that the simulated DMRs originate from the null hypothesis, they tend to be shorter and have smaller *Q*_*max*_. Consequently, the algorithm applies a range of thresholds for the minimum number of CpG sites and *Q*_*max*_, looking for a combination that would achieve the desired FDR level. If multiple sets of parameters achieve this FDR level, we select the one that filters out the fewest of the original DMRs.

#### Step III: Annotation

The final step of this part creates annotations of the final DMR list. Two types of annotations are currently implemented: associating DMRs with gene bodies and promoters, and with CpG islands. Users are required to provide the location data (gene list and CpG island list). These files are provided in https://carmelab.huji.ac.il/data.html for hg19. By default, RoAM defines the promoter region of each gene as 5,000 bp upstream of the transcription start site (TSS) to 1,000 bp downstream, but this can be set by the user. In addition, annotation can be done against any list of genomic segments, inserted as a BED file.

## Results

Epigenetics in general, and DNA methylation in particular, may respond to changes in internal or external conditions (23, 24). Research has unveiled connections between numerous environmental factors and alterations in DNA methylation (25–28). Consequently, even short bouts of environmental or lifestyle transitions may make epigenomic imprints that can be read.

Motivated by this, we decided to investigate potential epigenetic imprints of the Neolithic transition using RoAM. The Neolithic revolution is a pivotal milestone in human history, representing a significant shift in lifestyle, from primarily that of hunting and gathering to a more sedentary one based on agriculture and animal husbandry. Pre-Neolithic revolution humans typically lived nomadic life, relying on wild plants and animals for sustenance. The adoption of agriculture and domestication practices led to significant changes in diet, disease load, levels of physical activity and many other aspects of life. These changes might have been accompanied by biological and physiological changes (e.g. (29)). Emerging evidence suggests that DNA methylation in some genomic loci is sensitive to such lifestyle factors (30, 31). Hence, we decided to compare the ancient epigenomes of pre-Neolithic revolution to those of post-Neolithic revolution.

Ancient individuals sequenced to high coverage are still not abundant, and tend to represent very different populations. Comparing pre- to post-Neolithic revolution individuals across many different populations potentially adds confounding factors. To address this, we limited our study to 14 high-coverage individuals that come from the same region, the Balkans. Nine are pre-Neolithic revolution individuals, and five are post-Neolithic revolution ones (Table 1). Two samples, I1116 and I5725 are dated to a later period compared to the other post-Neolithic revolution samples. Whereas this chronological difference might introduce biases to the analysis, we nevertheless decided to include these samples in order to enlarge the number of post-Neolithic revolution individuals, which was anyway already small compared to the pre-Neolithic revolution group. Indeed, as will be shown below, these two samples sometimes show methylation patterns that are somewhat different than those of the other post-Neolithic revolution individuals. Genomic data of these ancient individuals were downloaded from the Allen Ancient Genome Diversity Project (32). Notably, petrous bone was the source for DNA extraction in all 14 samples, further reducing potential effects of confounding factors.

**Table 1.** List of samples used in this work. Sample names are taken from the Allen Ancient Genome Diversity Project.

| Sample name | Group | Sample Age (K years) | Location | Coverage |
| --- | --- | --- | --- | --- |
| I1507 | Pre-Neolithic revolution | 7.7 | Hungary (Tiszaszolos-Domaháza) | 22.42 |
| I4873 | Pre-Neolithic revolution | 7.9 | Serbia (Vlasak) | 25.76 |
| I4875 | Pre-Neolithic revolution | 8.5 | Serbia (Vlasak) | 21.48 |
| I4877 | Pre-Neolithic revolution | 8.5 | Serbia (Vlasak) | 27.44 |
| I4878 | Pre-Neolithic revolution | 7.8 | Serbia (Vlasak) | 25.3 |
| I4914 | Pre-Neolithic revolution | 8.1 | Serbia (Vlasak) | 24.85 |
| I5233 | Pre-Neolithic revolution | 8 | Serbia (Padina) | 23.55 |
| I5235 | Pre-Neolithic revolution | 10.8 | Serbia (Padina) | 25.61 |
| I5236 | Pre-Neolithic revolution | 10 | Serbia (Padina) | 26.91 |
| I1116 | Post-Neolithic revolution | 1 | Serbia (Gomolova) | 26.97 |
| I1496 | Post-Neolithic revolution | 7 | Hungary (Apc-Berekalya I) | 29.96 |
| I2520 | Post-Neolithic revolution | 5.1 | Bulgaria (Dzhulyunitsa) | 24.47 |
| I5077 | Post-Neolithic revolution | 7 | Croatia (Sopot) | 27.79 |
| I5725 | Post-Neolithic revolution | 2.5 | Croatia (Sv Kriz) | 27.49 |

We applied RoAM to reconstruct methylation for each sample and then detected DMRs between these two groups. Given the short time span separating the two groups, we expected to find only small methylation changes between them. We have therefore set the minimum methylation difference between groups (the Δ parameter, see Methods) to the very low value of 0.1.

Methylation patterns can exhibit significant variations between two distinct tissues within the same individual (33). As a result, much of the research in the field of paleoepigenetics has concentrated on the evolutionary aspects of the skeletal system (19, 34). The relevance of skeletal DMRs to changes in lifestyle following the Neolithic transition is debated. However, we have shown that there are loci in the genome where differential methylation in one tissue may reflect differential methylation in another tissue, as long as the methylation change occurs early during embryogenesis (4). To help in focusing on differential methylation that arose during such early developmental times, we crossed our results with published methylation data derived from blood samples of modern hunter-gatherers and farmers in Africa (35), where differentially methylated sites separating these population have been identified. We only considered genes that featured methylation changes in both bone and blood.

Our conservative analysis yielded only four DMRs between these pre- and post-Neolithic revolution populations (Table 2). Three of the DMRs are located inside gene bodies and, interestingly, all four overlap CpG islands, suggesting a possible regulatory role of these methylation changes. The DMR with the highest *Q*_*max*_ (406.4) was found inside the gene body of the PTPRN2 gene (also known as IA-2β, Figure 3A), which also harbors the third-highest number of differentially methylated sites in blood, separating modern African hunter-gatherers from farmers, with a total of 62 such sites. PTPRN2 is a transmembrane protein present in dense-core vesicles and represents a major auto antigen of type 1 diabetes (36). Previous works found that PTPRN2 has a key role in insulin secretion in response to glucose stimulus, and suppression or knocking down of this gene can impair this process (37–40). Epigenetic regulation on PTPRN2 has been examined previously, and a DNA methylation change in a CpG site within this gene, which does not overlap with the detected DMR, has been associated with childhood obesity (41).

**Table 2.** DMRs separating pre- and post-Neolithic revolution samples from the Balkan (ordered by *Q*_max_, from largest to smallest).

| Chrom. | DMR start (hg19) | DMR end (hg19) | Qmax | # CpGs | Gene | Pre-Neolithic revolution methylation | Post-Neolithic revolution methylation | Methylation difference |
| --- | --- | --- | --- | --- | --- | --- | --- | --- |
| 7 | 157405128 | 157407252 | 406.4 | 145 | PTPRN2 | 0.36 | 0.65 | 0.29 |
| 2 | 131008391 | 131011881 | 311.9 | 119 |  | 0.35 | 0.64 | 0.29 |
| 19 | 12983432 | 12985003 | 287.4 | 118 | MAST1 | 0.50 | 0.77 | 0.27 |
| 15 | 40266418 | 40269319 | 255.8 | 79 | EIF2AK4 | 0.40 | 0.74 | 0.34 |

**Figure 3.**
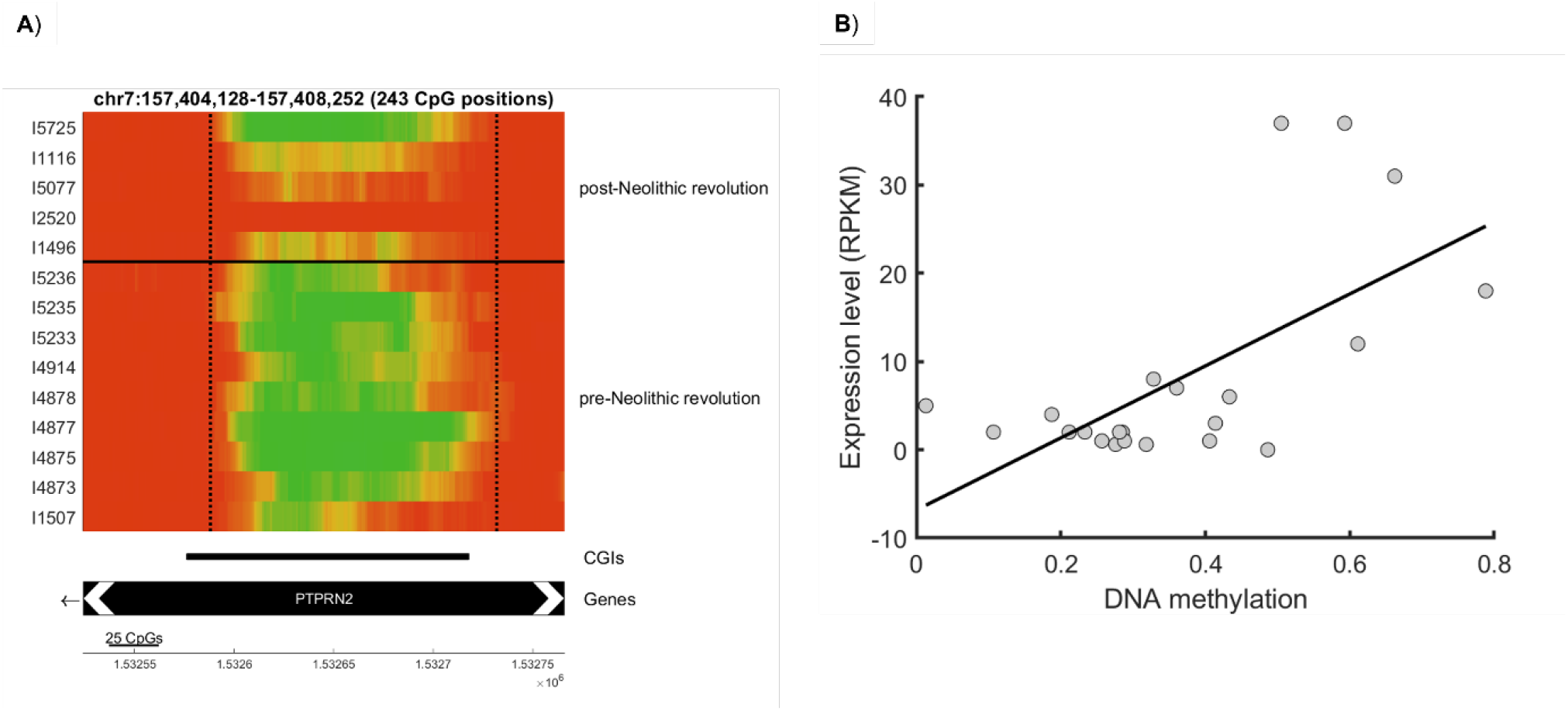
The DMR within the PTPRN2 gene. **A)** Reconstructed DNA methylation of the 14 Balkan samples. The DMR (dashed vertical lines) distinguishes between post-Neolithic revolution samples (upper lanes) and pre-Neolithic revolution ones (lower lanes). Methylation is color coded, from low methylation in green to high methylation in red. Lower lanes describe the genomic locations of CpG islands (CGIs) and genes. This DMR intersects a CpG island. **B)** Expression level of the PTPRN2 gene as a function of the mean methylation within the DMR, in 22 modern human tissues.

To find the predicted effects of the DNA methylation changes on the expression level of this gene, we looked at the correlation between the methylation in this DMR and the expression of PTPRN2 across 22 tissues of present-day humans taken from the Roadmap dataset (42). We found a significant positive correlation (*R* = 0.65, *P* = 8.8 · 10^−4^), suggesting that PTPRN2 was expressed in higher levels in post-Neolithic revolution individuals (Figure 3B). This implies lower insulin response to glucose stimulus in the pre-Neolithic revolution individuals.

Another DMR was detected inside EIF2AK4 (also known as GCN2, Figure 4A), a sensor for amino acid deprivation and a regulator of lipid metabolism and gluconeogenesis (43, 44). This kinase plays a crucial role in maintaining homeostasis during amino acid deprivation. When under a leucine-deprived diet, EIF2AK4 reduces insulin levels and increases insulin sensitivity (45, 46). However, in mice consuming a high fat diet, the opposite effect is shown, where EIF2AK increases blood insulin levels and decreases insulin sensitivity (47). Furthermore, EIF2AK4 is also implicated in diabetes, as its knockout in diabetic mice results in a decrease in serum fasting glucose and improved cardiac symptoms (48).

**Figure 4.**
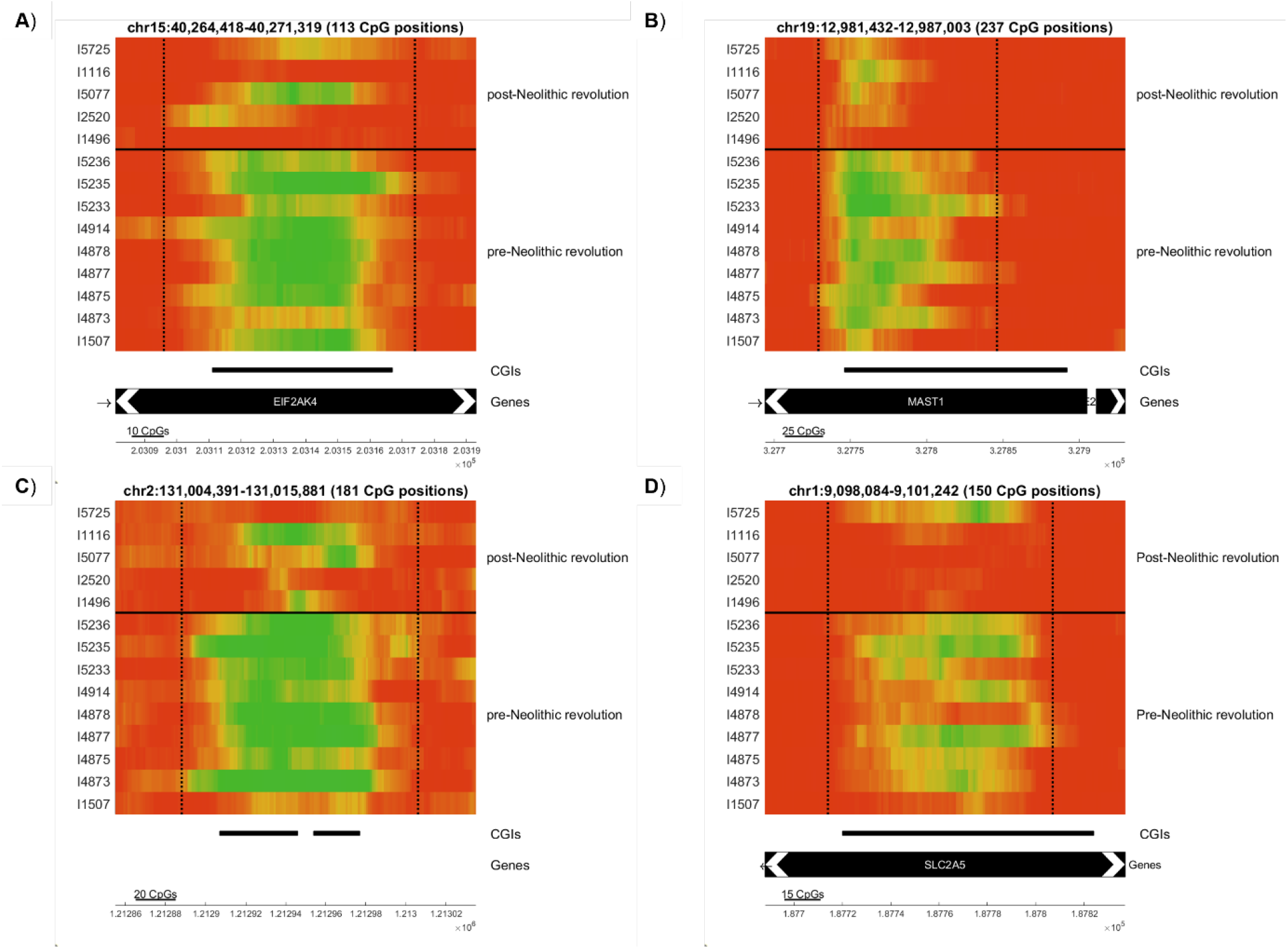
Additional DMRs (bounded by dashed vertical lines) that distinguish between post-Neolithic revolution samples (upper lanes) and pre-Neolithic revolution ones (lower lanes). Methylation is color coded, from low methylation in green to high methylation in red. Lower lanes describe the genomic locations of CpG islands (CGIs) and genes. **A-C)** The additional three DMRs detected in the analysis. **D)** The DMR with the highest *Q*_*max*_ that did not pass significance threshold. All DMRs intersect CpG islands.

The third DMR was detected in the gene MAST1 (Figure 4B), a kinase that plays a role in the central nervous system (49–51). No relation of this gene to metabolism or any function that might be related to the Neolithic revolution is currently known. The fourth DMR is not located on a promoter or a gene body (Figure 4C).

Only four DMRs out of an original set of 155,693 had properties that meet the thresholds set by the simulations to get an FDR below 0.05 (see Methods). Although not statistically significant, we noticed an interesting DMR that was very close to crossing the required thresholds. This DMR overlap CpG island, and is located within the SLC2A5 gene (also known as GLUT5, Figure 4D), a major fructose transporter in the gut. Many works showed that the presence of fructose stimulation can enhance, even within hours, SLC2A5 expression in the small intestine of adult animals. The same is true of neonatal and weaning pups that do not normally consume fructose and have low levels of SLC2A5 expression in their intestines (52–57). SLC2A5 is also associated with diabetes and obesity, as the gene is differentially expressed in insulin-sensitive tissues of patients with type 2 diabetes and in mouse models for diabetes and obesity, such as muscle (58) and fat tissues (59, 60).

## Discussion

We introduce here RoAM, a user-friendly program designed to provide a complete analysis pipeline for computational reconstruction of ancient methylomes and the identification of DMRs that distinguish ancient populations from each other. As the significance of evolutionary epigenetics is in the rise, RoAM proves to be a valuable tool for researchers seeking to integrate paleoepigenetic insights into their studies.

An advantage of RoAM is that new features are constantly added, gradually providing it with even more power. For example, a primary limitation of the reconstruction algorithm is that it cannot work on low-coverage samples, as the counts of Cs and Ts may be too low to allow for reasonable standard error of the estimator. To overcome this, we have introduced the concept of pooling, where counts from many low-coverage samples from the same group are amalgamated to provide a methylation map that represents the entire population (22). Pooling has already been integrated into RoAM, making it a viable tool for analyzing methylation maps in populations with low-coverage samples.

There are several limitations of the current software. First, the current implementation does not allow comparisons across more than two groups. Second, the code is limited to detecting DMRs between two groups that are exclusively composed of ancient samples. Ideally, we would like to integrate modern samples in the analyses, such that each group can potentially consist of a mixture of modern and ancient samples. Third, the algorithm exclusively performs methylation reconstruction on samples subjected to USER treatment (6). However, this treatment is not universally performed in aDNA library preparation. Finally, it does not account for difference in deamination rates along the read (6). We are currently actively working on developing solutions for all these limitations. RoAM will continue to be maintained and updated, with each solution promptly implemented in the code, providing RoAM with the capability to handle a growing number of samples of different types.

To showcase the algorithm, we conducted methylation reconstruction on 14 pre- and post-Neolithic revolution samples from the Balkans, and identified four DMRs that distinguish between them. The genes associated with these DMRs provide insights into understanding dietary changes that were induced by the Neolithic revolution. Two classic hypotheses claim that hypoinsulinism in pre-Neolithic revolution hunter gatherers provided an adaptive advantage. The carnivore connection hypothesis (61, 62) suggests that hunter-gatherers diet was high in protein and low in carbohydrates, and that in such conditions, insulin resistance would confer an evolutionary advantage, as it allows redirection of glucose to specific requirements such as embryonic development and brain functions. The thrifty genotype hypothesis (63) suggests that hypoinsulinism was a preferred strategy for storing food in times of food scarcity due to the instability in food sources. Glucose is specifically important for fetal development and to the function of the brain, and therefore hypoinsulinism can be a good adaptation to accommodate and supply the body needs when experiencing low or unstable glucose availability. In line with these claims, previous studies reported that hunter-gatherers from the north-western Kalahari display lower levels of blood insulin, while genetically similar communities that have adopted a sedentary lifestyle for 15 years show an increase in insulin levels during this period (64, 65). Additional work showed that short term consumption of a paleolithic diet can decrease insulin secretion (66). These works indicate hypoinsulinism in hunter gatherers, and specifically at lower insulin secretion. Our most pronounced DMR resides in PTPRN2, suggesting overexpression of this gene in post-Neolithic revolution individuals compared to pre-Neolithic revolution ones. Given its role in insulin secretion in response to glucose, this finding lends further credence to the claim that hunter-gatherers experienced hypoinsulinism. Further evidence for methylation changes in PTPRN2 that correlate with hunting and gathering lifestyle can be found in an independent study that compared methylation levels in modern hunter gatherers and genetically related farmers in Africa (35). In this study, PTPRN2 stand out as the gene with the third-highest number of differentially methylated sites, amounting to a total of 62 sites.

Another DMR lies with the EIF2AK4 gene. EIF2AK4 regulates insulin level and sensitivity in response to varied dietary components, and specifically during amino-acid deprivation. Changes in the expression level of this gene may be linked to the dietary shift during the Neolithic revolution. A plausible explanation for this change could be attributed to food scarcity, potentially resulting in the deprivation of certain amino acids for hunter-gatherers.

We also found a DMR within the SLC2A5 gene. This DMR is filtered out because of our very strict criteria, but it was just below the threshold, so we decided to discuss it here, as it might be potentially related to the Neolithic dietary transition. SLC2A5 is a fructose transporter, whose expression levels change when fructose consumption is increased. Our data do not allow us to determine the sign of the correlation between the methylation in this DMR and the gene expression, hence we cannot conclusively determine whether the methylation changes are associated with up or down regulation of this gene in post-Neolithic revolution times. However, we do observe a notable methylation change in this gene, that warrants further experimental examination.

It should be recognized that the DMRs we detect here likely represent just a small minority of the methylated changes that accompanied the Neolithic transition. First, we have used very strict filtering criteria, increasing precision on the expense of sensitivity. Second, the use of DNA methylation maps from bones means that we are able to identify only those methylation changes that occurred very early during embryogenesis, and simultaneously affect multiple tissues (4). We will not be able to observe tissue-specific methylation changes, where the methylation change is not shared with bone. We anticipate the existence of such tissue-specific methylation changes, particularly in genes associated with immune functions and metabolism.

## Acknowledgements

This publication was made possible through the support of a grant from the John Templeton Foundation (Grant ID# 61739 to L.C and E.M). The opinions expressed in this publication are those of the authors and do not necessarily reflect the views of the John Templeton Foundation. This study was also funded by the Israel Science Foundation [ISF grant 2436/22 to L.C. and B.Y.]. We wish to thank Shani Vaknine Treidel for help with the graphics.

L.C. is the Snyder Granadar chair in Genetics. E.M. is the Arthur Gutterman Family chair for Stem Cell Research.

